# Thermal stratification and fish thermal preference explain vertical eDNA distributions in lakes

**DOI:** 10.1101/2020.04.21.042820

**Authors:** Joanne E. Littlefair, Lee E. Hrenchuk, Paul J. Blanchfield, Michael D. Rennie, Melania E. Cristescu

## Abstract

Significant advances have been made towards surveying animal and plant communities using DNA isolated from environmental samples. Despite rapid progress, we lack a comprehensive understanding of the “ecology” of environmental DNA (eDNA), particularly its temporal and spatial distribution and how this is shaped by abiotic and biotic processes. Here, we tested how seasonal variation in thermal stratification and animal habitat preferences influence the distribution of eDNA in lakes. We sampled eDNA depth profiles of five dimictic lakes during both summer stratification and autumn turnover, each containing warm- and cool-water fishes as well as the cold-water stenotherm, lake trout (*Salvelinus namaycush*). Habitat use by lake trout was validated by acoustic telemetry and was significantly related to eDNA distribution during stratification. Fish eDNA became “stratified” into layers during summer months, reflecting lake stratification and the thermal niches of the species. During summer months, lake trout, which rarely ventured into shallow waters, could only be detected at the deepest layers of the lakes, whereas the eDNA of warm-water fishes was much more abundant above the thermocline. By contrast, during autumn lake turnover, the fish species assemblage as detected by eDNA was homogenous throughout the water column. These findings contribute to our overall understanding of the “ecology” of eDNA within lake ecosystems, illustrating how the strong interaction between seasonal thermal structure in lakes and thermal niches of species on very localised spatial scales influences our ability to detect species.

## Introduction

Environmental DNA (eDNA) is increasingly being used to conduct biodiversity surveys, species occupancy studies, and detect endangered and invasive species (Deiner et al., 2017; Taberlet, Coissac, Pompanon, Brochmann, & Willerslev, 2012). Molecular and bioinformatics techniques have become increasingly refined in order to optimise the capture of eDNA (Alberdi, Aizpurua, Gilbert, & Bohmann, 2017; Deiner, Walser, Mächler, & Altermatt, 2015), but much of the “ecology” of eDNA – its release, transport, distribution, and degradation – is still poorly understood (Deiner et al. 2017; Cristescu & Hebert, 2018). Recent studies suggest that the spatio-temporal distribution of eDNA in field settings is shaped by the seasonal dynamics of the system and behaviour of organisms (Bista et al., 2016; Handley et al., 2019), but these processes are generally understudied owing to the large spatial and/or temporal scales involved and the difficulty of obtaining high levels of biological replication at the habitat scale in order to make accurate inferences. Yet this knowledge is essential for adequate survey design and correct interpretation of results as we move into the genomic era of assessing eukaryotic biodiversity (Bohmann et al., 2014).

The spatial distribution of molecular signals within a habitat is shaped by both abiotic and biotic factors influencing the processes of shedding, persistence, transport, and degradation (Harrison, Sunday, & Rogers, 2019). Early eDNA studies examined the effects of single environmental factors on shedding and degradation in controlled environments such as aquaria or mesocosms, either with or without organisms present (Andruszkiewicz, Sassoubre, & Boehm, 2017; Klymus, Richter, Chapman, & Paukert, 2015; Lance et al., 2017; Mächler, Osathanunkul, & Altermatt, 2018). These studies were essential for determining the relative contributions to the distribution and persistence of eDNA particles. However, as eDNA matures into a tool that is being relied on for monitoring and environmental assessment, it is essential to understand the complex interplay between species’ habitat selection and spatio-temporal variation in abiotic factors in shaping the distribution of eDNA within ecosystems.

Abiotic factors such as temperature, water chemistry, and exposure to UV are thought to influence rates of eDNA shedding and/or degradation (Klymus et al., 2015; Lance et al., 2017; Sansom & Sassoubre, 2017; Sassoubre, Yamahara, Gardner, Block, & Boehm, 2016; Strickler, Fremier, & Goldberg, 2015). Abiotic factors also control eDNA transport at various scales in ecosystems, and therefore the spatial scale of species’ presence inferences. In aquatic ecosystems, speed and volume of lotic flow has received prominent attention in both experimental and field settings, with estimates of eDNA transport ranging from metres to kilometres (Deiner, Fronhofer, Mächler, & Altermatt, 2016; Jane et al., 2015). Similarly, studies in coastal marine waters demonstrate that although eDNA signals generally show decreasing community similarity at scales greater than 60-100 m, some signal transport still takes place, possibly as a result of particle transport by wave motion and water mixing (Donnell et al., 2017; Port et al., 2016).

By contrast, the influence of water movement on eDNA transport and species detection has largely been neglected for lacustrine systems. An important seasonal feature of many temperate lakes is stratification, where isolated layers of water are formed. During summer, the upper warm layer (epilimnion) is separated from a deep, cold layer of the lake (hypolimnion) by the formation of a thermocline (a temperature-dependent density gradient) between these layers. Brief periods of whole water-column mixing occur prior to and after stratification in dimictic lakes during spring and autumn (Wetzel, 2001). These hydrological layers give rise to distinct temperature and oxygen conditions that create different habitat niches for aquatic organisms. Thus, the seasonal cycle of lake stratification can concentrate organisms within, or isolate organisms from, certain habitats at different times of the year. The general view is that eDNA signal is more or less homogenous in freshwater lakes and ponds due to the relatively small size of such habitats when compared with the much larger and less discrete marine realm. However, there have been interesting insights from studies of single lakes which have found differences in eDNA community composition at the top and bottom of the water column, possibly indicating a role for the thermocline in separating these molecular signals (Hänfling et al., 2016).

Abundance, life history, physiology, and behaviour of organisms are implicated as biotic factors which shape the release of eDNA at varying scales. On a large geographic scale, the concentration of eDNA in water can reflect annual life history events such as migration or spawning, and can be used to track populations on the move or invasion fronts (Bylemans, Furlan, Gleeson, Hardy, & Duncan, 2018; Erickson et al., 2016; Spear, Groves, Williams, & Waits, 2015; Uchii, Doi, Yamanaka, & Minamoto, 2017). Several studies have used eDNA to monitor seasonal shifts in community assemblages in river estuaries (Stoeckle, Soboleva, & Charlop-Powers, 2017), coastal ecosystems (Berry et al., 2019; Sigsgaard et al., 2017), and large lakes (Bista et al., 2016; Handley et al., 2019). However, there have been few studies that look at within-habitat eDNA distribution particularly with respect to habitat niche specialisation or behavioural preferences (although see Macher & Leese, 2017; Nichols, Königsson, Danell, & Spong, 2012), and fewer still have examined how this might change seasonally. For some animals, habitat selection varies seasonally on relatively small spatial scales, but whether these changes are reflected in molecular signals remains largely unexplored.

Most freshwater organisms are ectothermic and optimize physiological performance by occupying habitats within specific thermal niches (Magnuson, Crowder, & Medvick, 1979). Thus, they have different thermal preferences according to their bioenergetic and foraging requirements. Many cold-water stenotherms, such as lake trout (*Salvelinus namaycush*), Coregonids, and sculpins (*Cottus* spp.) avoid the warm temperatures of lake surface waters during summer stratification due to the associated metabolic costs and increased oxygen requirements of doing so (Beitinger & Fitzpatrick, 1979; Ficke, Myrick, & Hansen, 2007; Magnuson et al., 1979). For example, lake trout display clear shifts away from littoral habitats when epilimnetic temperatures rise above 15 °C, suggesting that water temperature phenology is a strong determinant of seasonal habitat use (Guzzo, Blanchfield, & Rennie, 2017). In lakes where cold-water prey fish are absent, lake trout are known to make forays into the littoral zone in summer to access high-quality prey resources, although these trips are typically of short duration and constitute a small proportion of their total habitat use during warm summer days (Guzzo et al., 2017). Thus, habitat use by obligate cold-water species can be greatly reduced and constrained to deeper depths during summer stratification, especially in small temperate lakes where habitat volume reductions of >60% are common due to lack of preferred temperature and dissolved oxygen conditions (Paterson, Podemski, Wesson, & Dupuis, 2011; Plumb & Blanchfield, 2009). At the same time, opposite habitat restrictions would be occurring for warm-water fishes, resulting in the restriction of their distribution to the upper, warmer waters of lakes (McMeans et al., 2020).

Temperature-driven habitat segregation among species of freshwater fish has the potential to create depth-specific molecular signals during stratification. Temperate freshwater lakes often remain stratified for about half of the calendar year. Given that warm- and cold-water fishes spend most of their time at shallower and deeper depths respectively during stratification, it is likely that they release the bulk of their eDNA in these habitats. The general view is that eDNA signals of aquatic organisms are more or less homogenous in freshwater lakes and ponds, despite the distinct thermal preferences of the fish occupying these ecosystems. Thus, eDNA studies often involve the collection of surface water samples only, without considering the important seasonal forces which shape thermal stratification and the habitat preferences of organisms. However, there is emerging evidence that eDNA can reflect local species richness and also peak in concentration during seasonal events (Bylemans et al., 2018; Erickson, Merkes, & Mize, 2019; Harper, Anucha, Turnbull, Bean, & Leaver, 2018; Spear et al., 2015).

In this study we explored the impact of lake stratification and turnover on the distribution of eDNA in dimictic lakes and make specific predictions for warm- and cold-water fishes. We validated our results by simultaneously collecting detailed acoustic telemetry data to define fine-scale habitat preferences of an obligate cold-water stenothermic fish (lake trout). We hypothesised that: 1) Lake thermal stratification (i.e. summer) results in strong stratification of eDNA signals for species that are highly constrained (cold- and warm-water species) and less stratification for more generalist species (cool-water species) (Figure 1A). 2) Isothermal conditions (i.e. autumn turnover) result in homogenous eDNA signals for all thermal guilds of fishes throughout the water column (Figure 1B).

**Figure 1:**
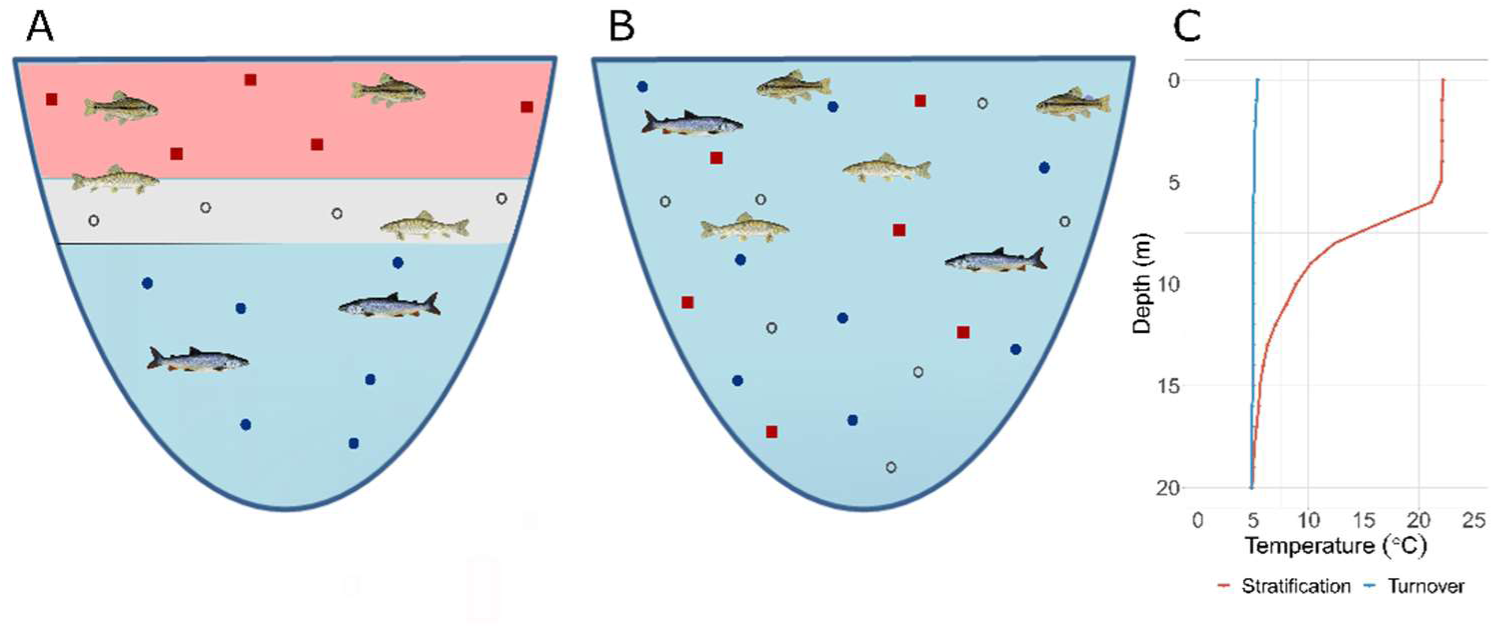
Conceptual figure showing hypothesised eDNA release in response to fish habitat selection and lake stratification/turnover. A lake during stratification (A) has isolated layers of water due to the formation of a temperature-dependent density gradient. There is minimal mixing between upper (epilimnion) and lower (hypolimnion) layers. Fishes select habitat due to bioenergetic requirements: this diagram shows potential habitat selection by warm-water, cool-water (able to inhabit all layers of the lake), and cold-water fishes. eDNA is released into stratified water layers and is slow to mix between the layers of the lake. Symbols represent the eDNA of warm-water fish (red squares), cool-water fish (open grey circles) and cold-water fish (filled dark blue circles). By contrast, during lake turnover (B) there is an isothermal water column with mixing between deep and shallow waters. Cold water fishes are now able to inhabit the entire water column. eDNA of all species is thoroughly mixed throughout the water column. Panel C shows temperature changes with lake depth during lake stratification (red line) and lake turnover (blue line) for Lake 373 during the 2018 sampling season.

## Materials and Methods

### Field collection

Sampling was conducted at the IISD Experimental Lakes Area (IISD-ELA), a remote research and monitoring facility in north-western Ontario, Canada. We sampled two lakes in summer and autumn of 2017 and repeated the summer and autumn sampling in five lakes in 2018. Study lakes vary in size from 25.8 - 56.1 hectares and have a maximum depth of 13.2 – 30.4m (Table S1). Monitoring of fish species at IISD-ELA has been conducted annually or bi-annually since the 1970s, therefore the species composition of most lakes is well known. There are 14 species of fish across all the study lakes (mean 8, range 6-10 species per lake, Table S2). All lakes have overlapping community compositions, including lake trout (*Salvelinus namaycush*), a cold-water top predator, in every lake. Sampling dates were chosen based on decades-long records of the timing of seasonal stratification and turnover (mixing) in these lakes. Moreover, temperature measurements of the water column were used to confirm lake stratification or turnover at the time of sampling (Table S3).

Water samples were taken at six depths, evenly dispersed throughout the water column at the deepest point of each lake (Table S3). Four 500 ml replicate water samples were taken per depth using an electrical pump and Jayflex PVC tubing (Winnipeg Johnston Plastics, MB, Canada) secured to a weight. To prevent contamination between lakes, dedicated tubing was used for each lake. Moreover, to prevent contamination among depth samples within a lake, the tubing was cleaned by flushing one litre of 30% bleach, then one litre of distilled water, followed by a two-minute flush of depth-specific lake water through the apparatus. For each sampling point, 500 ml of lake water was sampled and stored in an unused sterile Whirl-Pak (Nasco, ON, Canada) sealed within a large Ziplock bag. All samples were immediately transported to the lab in a cooler with ice packs and stored at 4 ⁰C until filtration. Water was filtered onto 47 mm 0.7μm pore GF/F filters using an electric vacuum pump and filtering manifold (Pall Corporation, ON, Canada). All filtrations were completed within eight hours of sample collection. One negative control of 500 ml distilled water was stored in the cooler and filtered in the same way as the field samples for each lake. The filters were immediately stored at −20 ⁰C and then shipped on dry ice to McGill University, Montréal for molecular analysis.

### Fish habitat use

We used published data on fish temperature preference to describe the thermal habitat use of fish species from the study lakes (Hasnain, Escobar, & Shuter, 2018; Hasnain, Shuter, & Minns, 2013, Table S2). For lake trout, we collected acoustic telemetry data on depth occupancy to determine seasonal habitat use and compared it with depth profiles collected with eDNA data. Extensive telemetry studies conducted at IISD-ELA over the past two decades have shown that the seasonal vertical distribution of lake trout is strongly influenced by prevailing temperature and oxygen conditions caused by stratification (Guzzo et al., 2017). Acoustic transmitter implantation into lake trout and data collection have previously been described in detail (Blanchfield, Flavelle, Hodge, & Orihel, 2005). Briefly, lake trout were captured by angling and surgically implanted with coded, acoustic, pressure-sensing telemetry tags (model V13P-1L; Vemco, Innovasea, Bedford, NS). Between 5 and 10 tagged lake trout adults were monitored in each lake during the study period. The pressure sensor on each tag was calibrated in the lake it was deployed in prior to implantation to ensure accurate depth readings (resolution: 0.08-0.15 m). The tags randomly emitted signals every 120-300 seconds (lakes 373, 626 and 239) or every 110-250 seconds (lakes 223 and 224). A number of data logging receivers (VR2W, 69 kHz; Vemco, Innovasea, Bedford, NS) were deployed under water at specific locations in the lake such that the “listening radius” of each receiver (spherical volume ~350 m diameter) overlapped slightly with the other receivers, resulting in maximum coverage of the lake. Each receiver was attached to a floating buoy and suspended ~2 m below the water’s surface or ~2-4 m above the bottom of the lake (dependent on mooring apparatus design). The receivers logged acoustic signals emitted by the tags through an omnidirectional hydrophone. Data (fish ID, date, time, pressure sensor reading) were continuously collected except when receivers were removed from the lake and downloaded (~8 h duration per lake, semi-annually). The pressure sensor data were converted to depth information using Vemco VUE software for each detection for the duration of the study (yielding ~200-700 depth detections for each fish in a typical 24-hr period). After downloading, duplicate detections (single tag signals detected by more than one receiver) were removed. In order to assess whether different time periods of cumulative eDNA persistence in the lakes affected the relationship between eDNA counts and telemetry data, we grouped telemetry data for each fish at different temporal scales, ranging from the day of eDNA sample collection, as well as one week, and one month prior to sample collection. The total number of detections of all fish were grouped into depth intervals reflecting the vertical distribution of the eDNA sampling (6 intervals per lake). We adjusted for varying depth interval size and variation in the total amount of telemetry detections for each lake over the relevant time period.

### Molecular analysis

DNA was extracted from filters using the Qiagen Blood and Tissue kit. We followed the manufacturer’s instructions with minor modifications: 370 µl buffer ATL was used in the initial incubation step, and the DNA was eluted in 2 x 60 µl of AE buffer. After elution, DNA was stored at −80 ⁰C. We included a DNA extraction control of blank sample for each lake. All samples were treated with the OneStep PCR Inhibitor Removal Kit (Zymo Research, Irvine, California). DNA was amplified in triplicate 12.5 µl reactions using 12S MiFish-U primers selected to target fish assemblages (Miya et al., 2015) tagged with Illumina adapters. The PCR chemistry was as follows: 7.4 µl nuclease free water (Qiagen), 1.25 µl 10X buffer (Genscript), 1 mM MgCl2 (ThermoFisher Scientific), 0.2 mM GeneDirex dNTPs, 0.05 mg bovine serum albumen (ThermoFisher Scientific), 0.25 mM each primer, 1U taq (Genscript) and 2 µl DNA in a final volume of 12.5 µl. PCR thermocycling followed a touchdown protocol with an annealing temperature from 66 to 64°C for 12 cycles followed by 28 cycles at 64 °C, which we found improved the proportion of samples which amplified. Negative PCR controls were included on each plate by substituting nuclease free water (Qiagen) for DNA. All filtration, extraction, and PCR negative controls were amplified in triplicate. PCR replicates from each sample were combined and cleaned with a 1: 0.875 ratio of AMPure beads. Samples were dual-indexed with v2 Nextera DNA indexes (Illumina). The samples were cleaned again with AMPure beads, quantified and equimolarised.

A DNA mock community of 27 North American fish species was constructed to evaluate the efficiency of our molecular methods and bioinformatics steps. DNA was extracted from individual fish samples using the Qiagen Blood and Tissue kit, following the manufacturer’s instructions, and equimolar DNA was combined to create the mock community. Two replicate libraries were PCR amplified and sequenced alongside the eDNA samples.

Libraries were sequenced over five lanes with either 91 or 92 samples in each. Sequencing was conducted using 2 x 250 bp Illumina MiSeq at Génome Québec, Montréal.

### Contamination prevention

Steps to prevent contamination were taken at each phase of work. During fieldwork, we used a dedicated boat and separate tubing for each lake to prevent between-lake transfer of DNA. All field equipment was decontaminated in 30% bleach and triple-washed with distilled water the evening before. Nitrile gloves were used when collecting the samples and changed between sampling points. The field lab used for filtering and storing of field equipment at IISD-ELA had not previously been used for sampling or storage of animal tissues. Benches were cleaned thoroughly with 20% bleach before use. After use, Buchner filtration funnels were washed in soapy water, soaked in 30% bleach for ten minutes, and vigorously triple-rinsed in ultrapure water between samples. DNA extraction and pre-PCR preparation were conducted in a dedicated environmental DNA lab at McGill University. The lab and equipment were thoroughly cleaned with 10% bleach before and after use (e.g. surfaces, floors, main shelving). Filter tips were used for all molecular work. There was no detectable PCR amplification in any field, DNA extraction or PCR negative controls based on gel electrophoresis, but we included all blanks for sequencing.

### Bioinformatics

We used custom scripts to remove adapters, merge paired sequences, check quality and generate amplicon sequencing variants (ASVs). Samples were received as demultiplexed fastq files from Génome Québec. Non-biological nucleotides were removed (primers, indices and adapters) using cutadapt (Martin, 2011). Paired reads were merged using PEAR (Zhang, Kobert, Flouri, & Stamatakis, 2014). Quality scores for sequences were analysed with FASTQC (Andrews, 2010). Amplicon sequencing variants (ASVs) were generated using the UNOISE3 package (Edgar, 2016), which uses a denoising pipeline to remove sequencing error and to cluster sequences into single variants (100% similarity). The generation of ASVs has several advantages over OTUs including finer resolution, accurate measures of diversity and easy comparison between independently processed datasets (Callahan, McMurdie, & Holmes, 2017). The full pipeline is available from https://github.com/CristescuLab/YAAP.

After ASVs were generated, we assigned taxonomy using BLAST+ (Camacho et al., 2009) and BASTA (Kahlke & Ralph, 2019), a last common ancestor algorithm (Supplementary Information). We used a custom reference database which contained only fish known to exist in the Lake of the Woods region (Ontario, CA), downloaded from NCBI. Biomonitoring has been ongoing since the 1960s so there is a well-developed knowledge of species composition in this area. We also compared our assignments against the full NCBI database and found only one additional fish ASV with the larger database. This matched to the *Hypophthalmichthys* genus (a carp), which is not known to exist at IISD-ELA but appeared at high abundance in one sample, possibly indicating a laboratory false positive. Other taxonomic groups appeared at very low frequencies when our ASVs were matched against the NCBI database, such as bacterial, mammalian and bird taxa, but as they were not the focus of our study they were excluded.

### Statistical approach

We used a variance stabilising transformation on our sample x ASV matrix to account for uneven library size across our samples. Unlike rarefaction, this approach does not discard valuable data due to differing library sizes (McMurdie & Holmes, 2014). We chose not to use a correction for the low numbers of sequences which appear in blank samples because we suspect that PCR amplification dynamics occur differently in samples which have extremely low amounts of template DNA when compared with positive template samples. Instead, information about sequences found in blank samples is displayed in Table S4. All statistical analyses were implemented in R v3.6.2 and vegan v2.5-6 (Oksanen et al., 2019; R Core Team, 2019).

We examined the relationship between fish community assemblages and the interaction between lake depth and lake state (stratified or isothermal) with PERMANOVA analysis. We used a Bray-Curtis distance matrix on our transformed sample x ASV matrix as the response variable. We tested the interaction between lake depth (coded as a continuous variable) and lake state on community composition, specifying 5000 permutations constrained within lake “strata”. We then tested for homogeneity in multivariate dispersion between our groups with the function betadisper. We used non-metric multi-dimensional scaling to visualise fish communities, by specifying either 2 or 3 dimensions (to minimise stress and achieve convergence) and 200 random starts.

We explored the contribution of each species to seasonal differences in ASV counts at different depths by fitting mixed effects models. We used ASV count for each species in each sample as the response variable modelled as the interaction between lake state (i.e. stratified or isothermal), depth of sample, and fish species to investigate whether stratification and turnover had variable effects for different species. We implemented negative binomial mixed effects models with lake identity as a random effect in glmmTMB (Brooks et al., 2017), using the total library size (DNA sequence counts for each sample) as a log offset in the model (Zurr, Ieno, Walker, Saveliev, & Smith, 2009). This approach allows us to control for library size while retaining interpretable response data (for example, in comparison to transforming variables which has been used in other studies). We also fitted several reduced models and compared these with AIC, always retaining the lake identity as a random effect term due to the nature of the experimental design. Once we had selected our best-fitting model with AIC, we confirmed the significance of the highest-level interaction term with a likelihood ratio test. Final models were evaluated for overdispersion.

We fitted a second series of mixed effects models to examine the relationship between the strength of eDNA signal in the water and habitat use by lake trout as detected by acoustic telemetry. We fitted the counts of lake trout ASVs as the response variable, and the interaction between lake state (stratified or isothermal) and telemetry detections as the explanatory variables, as this would allow the relationship to vary according to differential habitat use and presence of the thermocline. We implemented negative binomial mixed effects models with lake identity as a random effect in glmmTMB (Brooks et al., 2017), again using the total library size (DNA sequence counts for each sample) as a log offset in the model. This analysis was performed for each of the three temporal datasets of telemetry data collected (one day, one week and one month before the point of sampling), to test whether differences in the temporal range of habitat selection better explained the distribution of eDNA, as it is known to persist in the water column for several days to weeks. Several simpler models with a reduced fixed effects structure were fitted for each temporal dataset, and we compared all models with AIC.

## Results

### Thermal habitat structure

Temperature profiles in each lake confirmed that eDNA sampling occurred during stratification and turnover (isothermal or near-isothermal conditions) within the lakes under study (Table S3). The thermocline was confirmed as being between 4.60-6.60 m from the surface (approximately between eDNA sampling depths two and three for most lakes). These patterns are typical of those found in previous years during peak stratification and turnover for lakes in this region (Sichewski & Cruikshank, 1998).

### Recovery of eDNA sequences and taxonomic assignment

We recovered 94,013 ± 6,389 sequences per demultiplexed sample with an initial quality score of 33.0 ± 0.23. After removing adapters, discarding low quality sequences, merging paired end sequences and length filtering we retained 76,734 ± 5,954 sequences per sample. From the entire dataset we created 373 ASVs, onto which we were able to map back 98.6% of filtered sequences (Table S5). A total of 28 ASVs were assigned to fish species known to exist at IISD-ELA. Although this number was small as a proportion of the total number of ASVs, 95.1% of all the filtered sequences in the dataset belonged to fish found at IISD-ELA. The ASVs from other taxonomic groups had very low numbers of reads. This indicates that most sequences in our dataset belong to fish from this geographic region, rather than resulting from the amplification of non-target taxonomic groups (e.g. bacteria, birds and mammals, which appeared at very low frequencies, Figure S1).

In the mock community, we made 19/27 correct detections at species level. Of those not detected at species level, four were detected at genus level (i.e. the last common ancestor algorithm assigned a match of the correct genus with no species name), two were detected at family level (i.e. the correct family but no species or genus given by the last common ancestor algorithm), one had many congenerics detected although not the correct species, and one could not be detected at any level.

The eDNA samples detected the majority (12/14) of fish species confirmed by both historical and present-day fishing surveys as being present in these habitats. The two species which were not detected (brook stickleback and longnose dace) are known to prefer near-shore and stream habitats and are also noted as being rare in many of these lakes, and thus sampling at the centre point of the lake may not be optimal to detect them at these times of year. We were able to assign the majority of ASV sequences at species-level using the last common ancestor algorithm with two exceptions. *Coregonus artedi* could only be assigned at genus level, as a closely related congener (*Coregonus clupeaformis*) also exists in this region (although *C. clupeaformis* is not present in any of our study lakes). *Chrosomus neogaeus* and *Chrosomus eos* were both assigned at genus level, possibly because pure *Chrosomus eos* does not exist in this region but instead forms both cytoplasmic and nuclear hybrids with *Chrosomus neogaeus* (Mee & Taylor, 2012).

### Fish community assemblages

During stratification, the relative proportions of ASVs from each species per sample changed dramatically at different depths in the lakes (Figure 3A). The overall species composition of the lakes was the same, yet species detection differed greatly at certain depths; with the greatest change taking place between points 2 and 3, which demarcates the thermocline in most lakes. For example, eDNA from cold-water stenotherms could only be detected in large proportions at the bottom of the lakes during lake stratification (lake trout *Salvelinus namaycush* and slimy sculpin *Cottus cognatus*). Lake trout were not detectable at all at the shallowest measurement points (1-1.5 m from the surface) at this time. Warm-water minnow species, which habitually inhabit shallow and littoral waters (e.g. *Chrosomus neogaeus, Margariscus margarita*, and *Pimephales promelas*), were detected in much greater proportions at the surface, with large decreases in the proportions of sequences in samples taken from below the thermocline. eDNA from cool-water eurytherms was distributed across all sampling depths, with the exception of *Coregonus*, which was only abundant at points 2 and 3 and could barely be detected at either the shallowest or deepest depths.

During lake turnover in late autumn, fish community detection by eDNA was much more homogenous throughout the different depths of the lake (Figure 3B), characterized by a greater proportion of cold-water fish sequences found at shallow depths. Changes in detection throughout the water columns were relatively small; for example, there was a slight increase in the proportion of *Cottus cognatus* sequences recovered at deeper sampling depths, but this species was found in the shallow samples as well. Similarly, there was a slight decrease in the sequences of minnow and perch species at deeper depths in the water column (*Perca flavescens, Margariscus margarita, Pimephales promelas*), but minnows could still be detected at the deepest depths in greater proportions than during stratification. *Coregonus* detections were no longer concentrated to the middle of the water column but could be detected at shallow and deep depths as well.

There was a significant interaction between lake depth and lake state affecting fish community assemblages detected by eDNA (PERMANOVA, F_1,335_ = 4.35, p = 0.0002). This result indicates that fish communities were detected throughout the water column differently if the lake was stratified or isothermal. NMDS plots for each lake showed that communities were clearly grouped by lake state (Figure 2), with distinct communities detected during stratification and turnover in most lakes. This result was confirmed by our mixed effects modelling approach to describe the distribution of fish ASV counts. The model which best fit the data included the three-way interaction between lake state (stratified or isothermal), eDNA sample depth, and fish species as an explanatory factor, when compared to any reduced model (ΔAIC 92.7). A full list of the reduced models that we tested and their AIC scores appears in Table S6. The three-way interaction between lake state, sample depth and species was highly significant (likelihood ratio test = 112.7, p < 0.001). eDNA from different fish species was distributed across the vertical column differently in each water mixing period.

**Figure 2:**
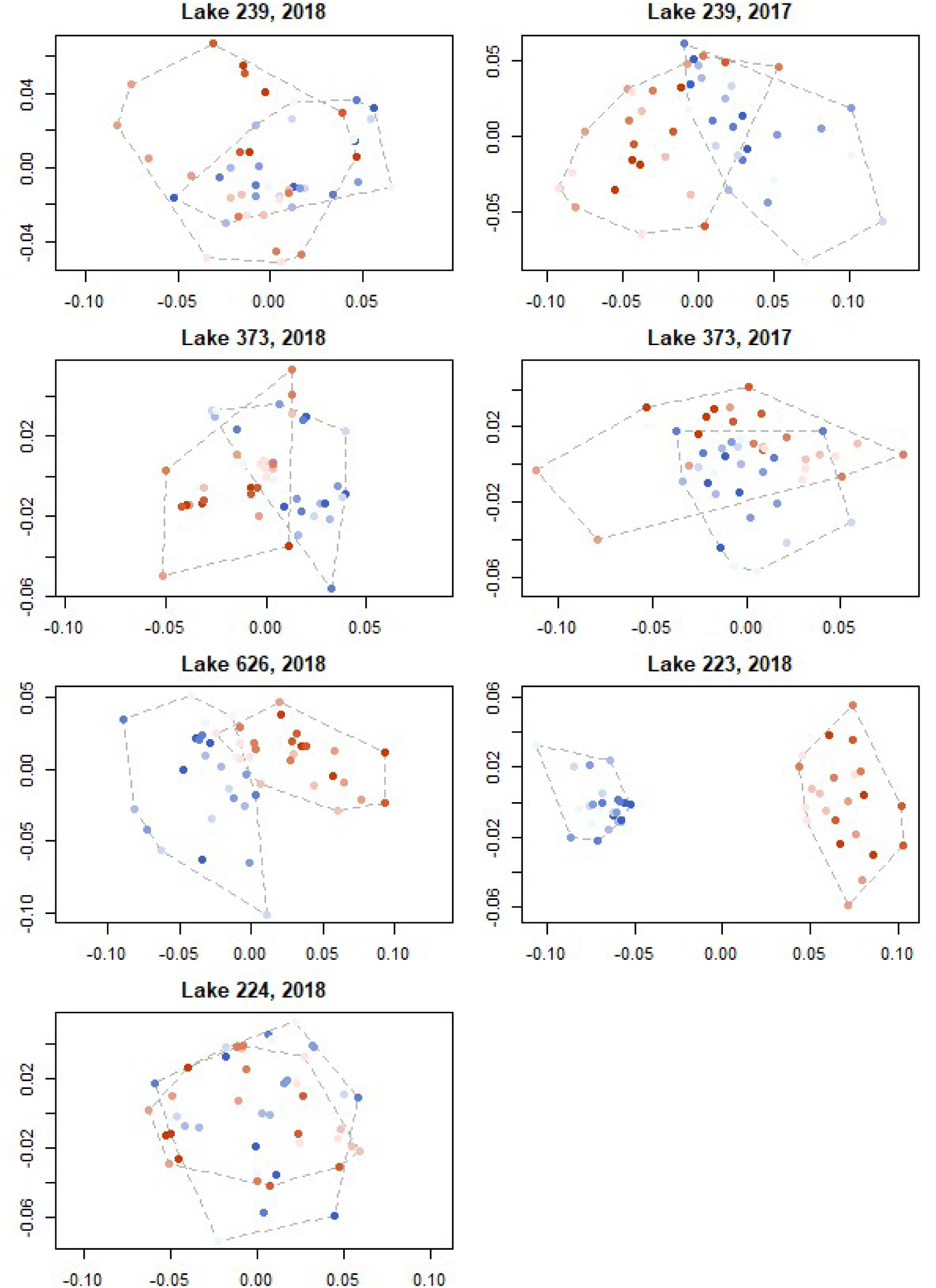
NMDS plots for each lake showing community dissimilarities detected by each sample. Samples from different seasonal water conditions are coloured differently (stratified samples in red, turnover samples in blue). The intensity of colour varies according to sample depth in the water column: the shallowest samples are represented with the lightest colours and the deepest samples with the darkest colours.

### Relationship between eDNA and lake trout habitat use

Lake trout eDNA was primarily concentrated in the bottom half of lakes (Figure 4A) during lake stratification (corresponding to points deeper than 6.25 - 16.5 m depending on the lake sampled). During lake turnover, lake trout eDNA was very abundant at all points in the water column, with no clear patterns according to sampling depth. Acoustic telemetry showed the lake trout inhabited the bottom two thirds of the water column during stratification, although they were less likely to occupy the deepest depths (Figure 4B red bars, median depth of telemetry detections = 7.74 - 11.90 m). During turnover, lake trout primarily selected habitat in the top third of the water column, with frequency tailing off at the deepest part of the lake (Figure 4B blue bars, median depth of telemetry detections = 1.73 – 6.51 m). The difference between median depths of fish one month and one week before, as well as the day of sampling was not large (Table S7).

The top ranked model to explain lake trout eDNA counts included the interaction between lake state (stratified or isothermal) and telemetry detection frequency for the month prior to the day of sampling (log(*lake trout ASV counts*) = −2.14 + 6.80*telemetry* + 0.97*turnover* - 6.02*telemetry x turnover*). There was a positive correlation between lake trout telemetry detections and eDNA counts during lake stratification, but no relationship during turnover (Figure 5). There were also five other models within two AIC counts of the top ranked model, which could be considered as having equal explanatory power (all models are listed in Table S8). These included a model with only the two main effects (no interaction) for average telemetry detections for the data from a month prior to sampling, as well as models with and without the interaction term for both the week prior to sampling, as well as the day of sampling, indicating that there were not large differences in the abilities of the different temporal groupings of telemetry detections to predict lake trout eDNA.

## Discussion

Our study was designed to test the influences of lake stratification and mixing on eDNA distribution within the framework of a replicated, whole-lake experimental design. Our results demonstrate that eDNA signals show very strong seasonal stratification during summer and mixing during autumn in a manner that closely reflects the thermal preference of fishes. We detected large differences in fish community composition during different lake states (Figure 2). During stratification, the most dramatic changes in community composition measured with eDNA took place in samples above and below the thermocline: warm-water fish eDNA was stratified above the thermocline and cold-water fish eDNA was concentrated below the thermocline (Figure 3). These differences were observed even across very small spatial scales (<30 m) between shallow and deep sampling points. By contrast, during lake turnover, eDNA of all fish species was relatively homogenous throughout the water column.

**Figure 3:**
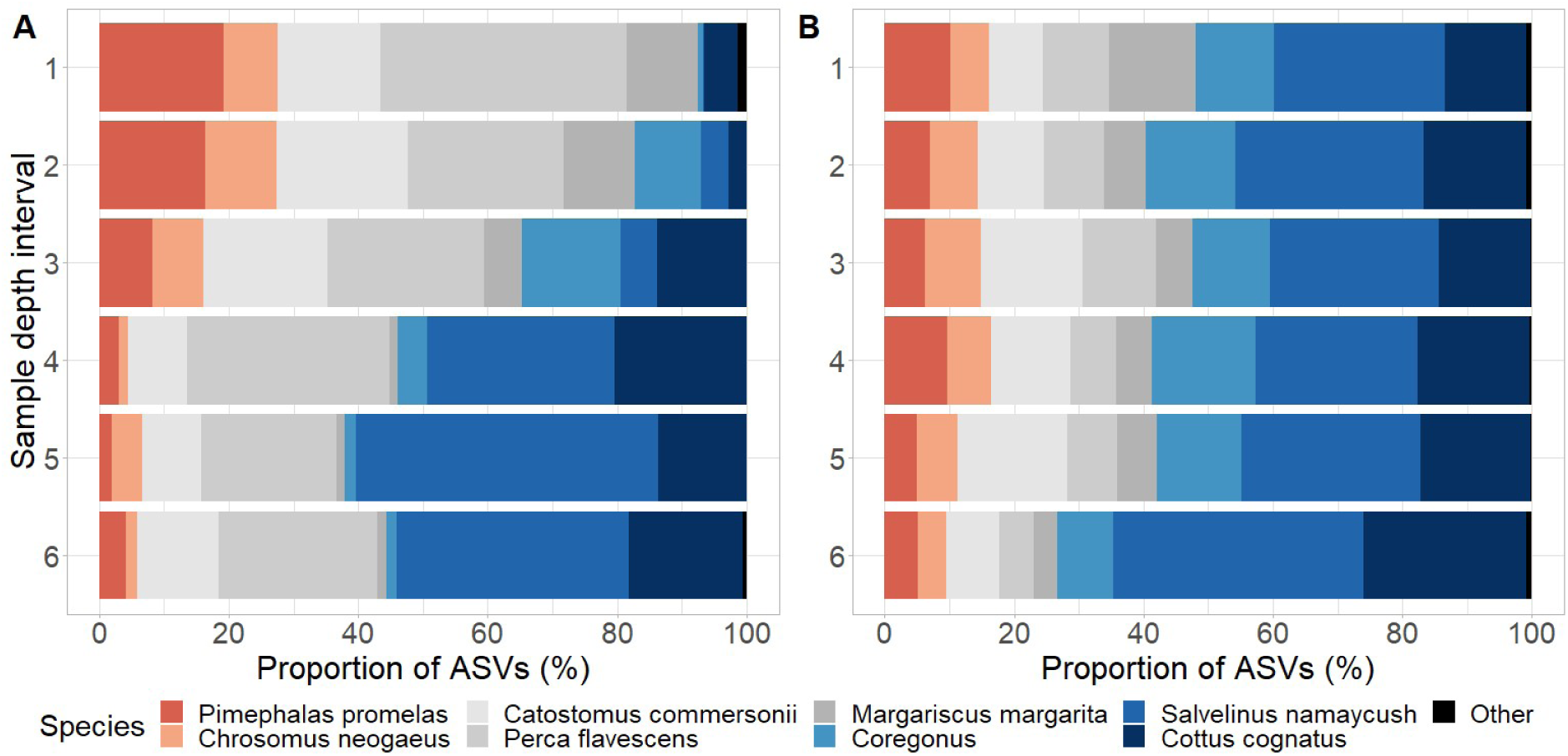
Proportional barplot shows the relative species composition detected by amplicon sequencing variants (ASVs) of all lakes combined during lake stratification (A) and lake turnover (B), at different sample intervals in the water column. The depth variable comprises of six evenly spaced vertical sampling points in the water column, and thus absolute measurements will vary for lakes of different depths. Point 1 is the shallowest measurement near the surface of the lake. Fish species are arranged in order of warm to cold thermal guilds (Table S2).

Few studies have managed to weigh the relative importance of abiotic and biotic influences on the distribution of eDNA – in this system, the two are intrinsically linked through bioenergetic requirements of fish which are manifest as thermal preferences. Thermal density gradients of lake water during stratification create distinct microhabitats for lake trout that provide suitable oxythermal habitat, which is generally defined as the volume of the lake that is <15°C with >4 mg L^−1^ DO (Plumb & Blanchfield, 2009). In late summer, optimal oxythermal habitat for lake trout is greatly reduced, concentrating this species into a narrow band within lakes that is often only a few meters thick (Plumb & Blanchfield, 2009). As a result, lake trout eDNA becomes localised due to narrow habitat selection by this cold-water stenotherm and the presence of the thermocline, which restricts water mixing between the epilimnion and hypolimnion (Wetzel, 2001). This is an important finding for the design of eDNA sampling studies, given that our study lakes are some of the smallest capable of supporting lake trout habitat. During lake turnover, the shallow-water presence of lake trout (shown by acoustic telemetry results to be in the top third of the water column) is decoupled from the distribution of eDNA signals, highlighting the role that water column mixing may have to play in dispersing the eDNA signal (Figure 4). Rapid cooling of epilimnetic waters in autumn initiates complete water column mixing and at the same time triggers lake trout movements from the hypolimnion to the shallow littoral areas of the lake to spawn in early-mid October. These abiotic and biotic processes result in a large amount of eDNA redistribution and release, respectively, leading to relatively even eDNA distribution throughout the water column.

**Figure 4:**
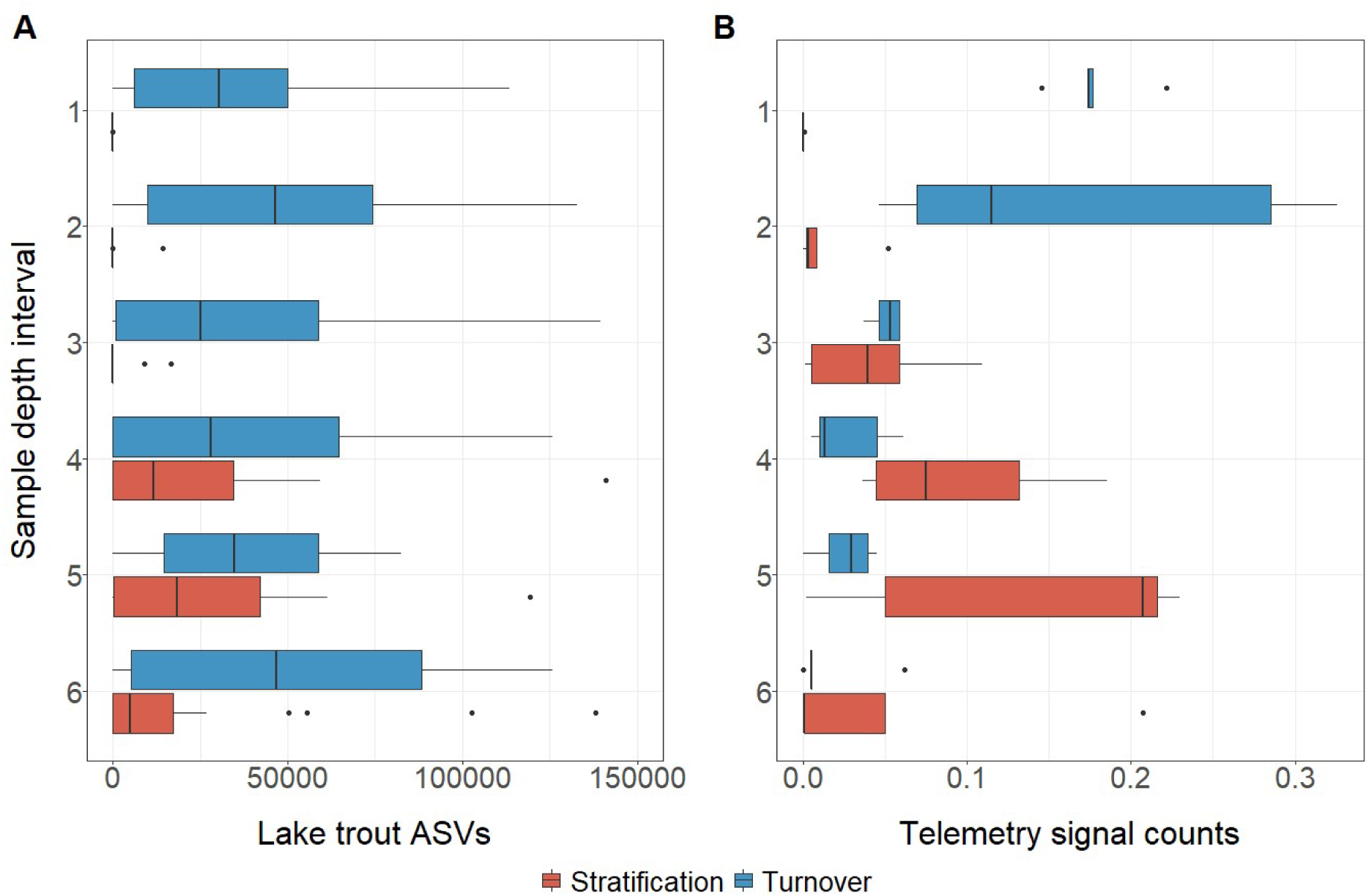
Lake trout amplicon sequencing variants (A) and lake trout telemetry detections (B) ordered by lake depth with stratified samples in red, turnover samples in blue. The depth variable is comprised of six evenly spaced vertical sampling points in the water column, and thus absolute measurements will vary for lakes of different depths (minimum lake depth = 13.2 m, maximum lake depth = 30.4 m). Point 1 is the shallowest measurement near the surface of the lake. Telemetry signal counts are expressed as a proportion of the total telemetry counts for that lake over the previous month. Depth interval size is also controlled for.

**Figure 5:**
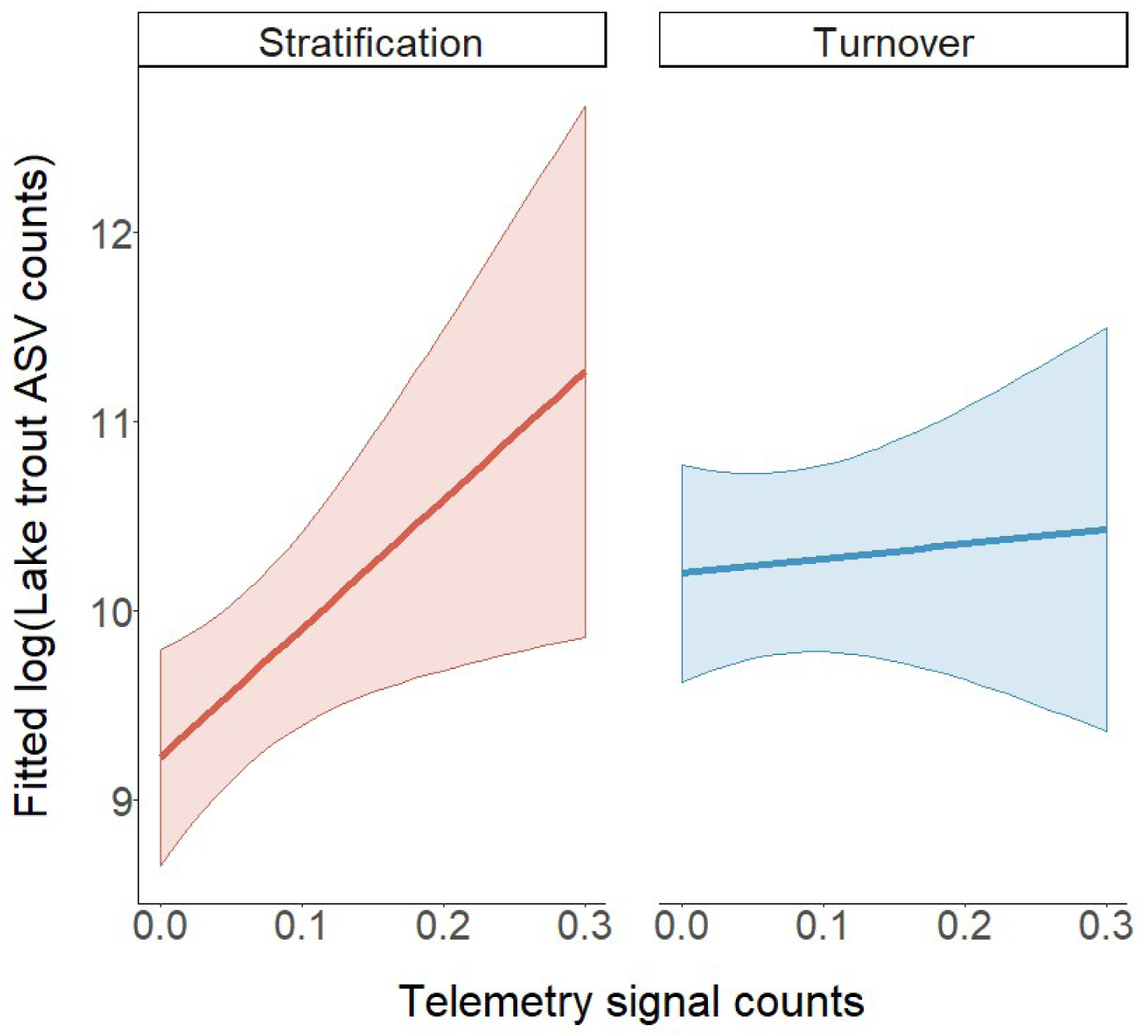
Model predictions from the best fit model to explain lake trout amplicon sequencing variants (ASVs). The best fit model included the interaction between seasonal water column thermal structure and proportion of telemetry signals in that depth interval. Telemetry signal counts are expressed as a proportion of the total telemetry counts for that lake over the previous month, and depth interval size is also controlled for. Shaded error bars are 95% confidence intervals.

Results from other fish species also suggest the importance of lake state (stratified or isothermal) in isolating or dispersing eDNA signals in lacustrine systems after initial eDNA release. The creation of microhabitats according to temperature gradients resulted in the detection of distinct community assemblages above and below the thermocline. During stratification, large amounts of eDNA from warm-water minnow species such as *Pimephales promelas* and *Chrosomus neogaeus* were found at the shallowest depths of the lake (the shallowest two sampling points fell between 1 and 6.25 m), consistent with their observed association with littoral regions of IISD-ELA lakes (Guzzo et al. 2014), and documented temperature preferences (Table S2). Moreover, eDNA sampling during lake turnover showed a much more equitable distribution of eDNA signals for warm-water minnow species. Thus, the contribution of water mixing to transporting warm-water fish eDNA to the bottom of the lake and shaping the distribution of eDNA is likely to be considerable. Interestingly, the minnows in our study lakes are classified as littoral-benthic species, spending the majority of time at the shoreline and small streams around the edges of the lake, indicating that the water between the shoreline and centre point in the epilimnion is well mixed. Studies involving the addition of tritiated water to the epilimnion of dimictic lakes have confirmed that the composition of the epilimnion becomes homogeneous one day after tracer injection, with vigorous mixing primarily occurring due to wind-induced horizontal movement. By contrast, rates of vertical diffusion of tracer across the thermocline of stratified lakes are much slower (Quay, 1980). Few studies have considered how habitat selection by organisms shapes the release of their eDNA or how this should influence design of biomonitoring surveys with eDNA.

Around the world, lake habitats have a variety of mixing regimes and other water movements which could influence the distribution of eDNA. Stratification is a major structuring force in temperate lakes, as long as the lakes are deep enough to allow for the formation of a thermocline. Potentially, deeper lakes will have more distinct microhabitat isolation between the epilimnion and deep waters, which in turn might result in a greater isolation of warm-water and cold-water species’ eDNA above and below the thermocline. Our results reflect those of Handley et al., (2019), who found greater heterogeneity in community composition of samples at three depth points during summer sampling when compared with winter sampling in their study of a single deep lake (1480 ha, depth of 44m/64m in two basins), and that eDNA from a cold-water stenotherm (*Salvelinus alpinus*, Arctic char) was only detectable in midwater and deep water habitats. Such findings may also apply to other monomictic, dimictic and meromictic lakes, as well as tropical and temperate oceans, which undergo periods of seasonal or permanent stratification. By contrast, Li et al., (2019) found eDNA of deep water species in shoreline samples during winter sampling, but as it is not clear to what degree (if any) the study lakes are stratified during winter months, this may have been the result of thorough mixing during autumn turnover. While previous eDNA studies have highlighted the surprising potential of rivers and streams to transport eDNA in the range of hundreds of metres to kilometres (Deiner & Altermatt, 2014; Deiner et al., 2016; Jane et al., 2015); we show that other hydrological forces can isolate microhabitats from each other which are physically just a few metres apart.

As with all ecological sampling techniques, there are a number of potential routes for false positives and negatives to occur with eDNA sampling in the field (Ficetola et al., 2015; Jerde, 2019). Increased biological and technical sampling effort, coupled with adequate preservation of DNA has already been called for to limit false negatives (Ficetola et al., 2015), but it is apparent from our analysis that carefully planning the timing of sampling and/or location of samples is highly important, when a difference of even a few metres could alter conclusions regarding species presence or absence. Maintaining the status quo of a surface sampling approach during the summer months will exclude or limit the consistent detection of cold-water species during periods of seasonal stratification, resulting in poor representation of these species in datasets. By planning monitoring campaigns for lake turnover, practitioners can use surface samples (which are often easier and faster to collect) to reliably sample fish species with a wide range of bioenergetic requirements. If sampling must be carried out during lake stratification, cold-water species can be targeted by sampling deeper layers with pumps, Freidinger/van Dorn bottles or integrated samplers (e.g. Handley et al., 2019; Hänfling et al., 2016; Lim et al., 2016; Yamamoto, Masuda, Sato, Sado, & Ara, 2017), as well as sampling surface waters to detect eurytherms. Use of this equipment presents further challenges in the field if sampling of multiple habitats is planned, as careful cleaning of equipment between habitats is necessary to reduce cross-contamination.

Much advancement has been made in molecular and computational approaches for eDNA work, confirming methods of substrate filtration, DNA extraction, primer choice, and bioinformatic filtering (e.g. Alberdi et al., 2017; Clare, Chain, Littlefair, & Cristescu, 2016; Deiner, Walser, et al., 2015). The design of field sampling campaigns provides the foundation on which other methods build, including timing and duration of sampling, location and replication of samples, power of experimental design, and even choice of sampling equipment. Many early studies used mesocosm approaches to study the fieldwork components of eDNA work, such as the abiotic and biotic influences on the rates of DNA production and degradation (e.g. Mächler et al., 2018; Seymour et al., 2018; Strickler et al., 2015). Using this approach, environmental factors can either be studied in isolation or as a multifactorial experiment in combination with a low number of other variables, while allowing for experimental replication and some control of other sources of environmental variation. Yet, there are many interacting facets that control the rates of production, transport and decay of eDNA within ecosystems that cannot be observed within small artificial systems, as has been argued in other areas of ecology which make use of mesocosm studies (Carpenter, 1996). Equally, the ecological significance of these factors cannot be tested when examined in isolation (Carpenter, Chisholm, Krebs, Schindler, & Wright, 1995).

Studies at the habitat scale have already suggested possible generalities linking eDNA to biological activity; for example, that peaks of eDNA can indicate the onset of reproduction (Bylemans et al., 2018; Spear et al., 2015), or relative abundance of species (Li et al., 2019). Our next challenge in eDNA research will be to scale up experimentation to produce generalisable rules for eDNA distribution in real ecosystems and interpret this in light of the biology of our study organisms.

## Animal care permits

Fish were collected and the telemetry tags implanted under the following permits: Ontario Ministry of Natural Resources and Forestry Licence to Collect Fish for Scientific Purposes #1085769 (2017), #1089495 (2018) and Lakehead University Animal Use Protocol #1464657 (renewed in 2017 and 2018).

## Acknowledgements

We are indebted to the many IISD Experimental Lakes Area students and staff for maintaining records of field data and for logistical assistance with this project. S Michaleski, R Henderson, P Bulloch, M Haust, C Jackson and A McLeod contributed specific field assistance to this project. K Sandilands provided equipment used in this project. This work was funded by a Mitacs Accelerate Industrial Fellowship (JEL), an NSERC Collaborative Research and Development award (MEC), Canada Research Chair and NSERC Discovery award to MEC and MDR, Québec Centre for Biodiversity Science Excellence award (JEL), the WSP Montréal Environment Department and in-kind support from the IISD Experimental Lakes Area and Fisheries & Oceans Canada. We thank Jean Carreau and Patrick LaFrance of WSP Montréal for useful discussions on the topics of eDNA and biomonitoring.

## Data Accessibility

Raw fastq files and the sample x ASV table are available at Dryad (data to be uploaded upon manuscript acceptance). Scripts to process bioinformatic data are available from https://github.com/CristescuLab/YAAP.

## Author contributions

JEL and MEC designed the experiments. JEL and LEH collected molecular data, JEL analysed the experiments and made the figures. LEH, PB and MR collected and processed the telemetry datasets and ongoing surveys of species richness. JEL wrote the first draft of the manuscript and all authors contributed to editing.

