## Supplementary material for "Thermal stratification and fish thermal preference explain vertical eDNA distributions in lakes"

### Table of contents


| Item | Page number |
|---|---|
| Table S1: Physical properties and population characteristics of each lake | 2 |
| Table S2: Species composition in the lakes and thermal guild classification | 3 |
| Table S3: Temperature/depth profile for each lake. | 4 |
| Table S4: Sequences assigned in blank samples | 5 |
| Table S5: Summary of sequences retained at each stage of bioinformatic analysis | 6 |
| Table S6: Candidate model structures and AIC values to assess species Amplicon Sequence Variant (ASV) count as a function of sampling depth, lake state, and fish species | 7 |
| Table S7: Average depths of lake trout during stratification and turnover | 8 |
| Table S8: Candidate model structures to assess lake trout Amplicon Sequence Variant (ASV) count as a function of proportion of telemetry detections and lake state | 9 |
| Figure S1: Numbers of taxonomically assigned ASVs using a local copy of the NCBI database | 10 |

Table S1: Physical properties and population characteristics of each lake at IISD Experimental Lakes Area. We selected lakes which contain lake trout and typically experience stratification during the summer months.

| Lake | Coordinates | Trout population size (lcl, ucl) | Size (ha) | Maximum depth (m) | Inflows | Outflows | Sampling months |
|---|---|---|---|---|---|---|---|
| **ELA 626** | N49.75279<br>W093.79799 | 242<br>(207 - 282) | 25.8 | 13.2 | Headwater lake | 1 | August 2018<br>October 2018 |
| **ELA 239** | N49.39493<br>W093.43211 | 70<br>(55 - 90) | 56.1 | 30.4 | 3 | 1 | July 2017<br>October 2017<br>August 2018<br>October 2018 |
| **ELA 373** | N49.44422<br>W093.47589 | 345<br>(316 -377) | 28.0 | 21.5 | Headwater lake | 1 | July 2017<br>October 2017<br>August 2018<br>October 2018 |
| **ELA 224** | N49.69000<br>W093.71694 | 387<br>(367 – 409) | 25.9 | 27.4 | 1 | 1 | August 2018<br>October 2018 |
| **ELA 223** | N49.698333<br>W093.70778 | 437<br>(279 - 683) | 27.3 | 14.4 | 1 | 1 | August 2018<br>October 2018 |

Table S2: Species composition in the lakes and thermal guild classification

We present presence/absence and final temperature preference (FTP) for all fish species in our study. FTP is the temperature that fish gravitate towards when provided with a broad range of temperatures.

| | **Lake** | | | | | **Temperature metrics from [1], [2]** | |
|---|---|---|---|---|---|---|---|
| **Species** | **223** | **224** | **239** | **373** | **626** | **Thermal guild** | **FTP** |
| *Couesius plumbeus* | x | | | | x | WARM | 27.0 |
| *Pimephales promelas* | x | x | | | x | WARM | 26.6 |
| *Chrosomus eos* | | x | | x | | WARM | 25.3 |
| *Chrosomus neogaeus* | x | x | x | x | x | COOL | 24.1 |
| *Catostomus commersonii* | x | x | x | x | x | COOL | 23.4 |
| *Etheostoma exile** | | | x | | | COOL | 22.8 |
| *Culaea inconstans* | x | x | | | | COOL | 21.3 |
| *Esox lucius* | | | x | | | COOL | 20.7 |
| *Perca flavescens* | | | x | | x | COOL | 17.6 |
| *Margariscus margarita*** | x | x | | x | x | COOL | 16.0 |
| *Rhinichthys cataractae* | | | | | x | COOL | 15.3 |
| *Coregonus artedi* | | | x | | | COLD | 12.4 |
| *Salvelinus namaycush* | x | x | x | x | x | COLD | 11.8 |
| *Cottus cognatus* | x | x | x | x | x | COLD | 11.0 |

* Data not available, data for Johnny Darter used.

** Data from Stauffer et al. 1984 used [3]

Table S3: Temperature/depth profile for each lake for two lake states, stratification (July/August) and turnover (October).

Intervals containing an eDNA sampling point have been highlighted in bold.

| **Depth (m)** | **626 Aug 2018** | **626 Oct 2018** | **239 Aug 2018** | **239 Oct 2018** | **224 Aug 2018** | **224 Oct 2018** | **223 Aug 2018** | **223 Oct 2018** | **373 Aug 2018** | **373 Oct 2018** |
|---|---|---|---|---|---|---|---|---|---|---|
| | Temperature measured in °C | | | | | | | | | |
| 0.0 | 22.4 | 4.6 | 22.7 | 4.4 | 22.5 | 5.3 | 23.1 | 4.8 | 22.2 | 5.4 |
| 1.0 | **21.7** | **4.9** | **22.6** | **4.5** | **23.0** | **5.4** | **23.0** | **4.8** | **22.1** | **5.3** |
| 2.0 | **21.6** | **5.0** | 22.6 | 4.5 | 23.1 | 5.5 | 22.8 | 4.8 | 22.1 | 5.2 |
| 3.0 | 21.6 | 5.1 | 22.5 | 4.5 | 23.1 | 5.5 | **22.8** | **4.8** | 22.1 | 5.1 |
| 4.0 | **21.5** | **5.1** | 22.5 | 4.5 | 23.0 | 5.5 | 22.7 | 4.7 | **22.1** | **5.1** |
| 5.0 | 21.1 | 5.1 | 19.6 | 4.5 | **22.8** | **5.5** | **21.5** | **4.7** | 22.0 | 5.1 |
| 6.0 | **16.6** | **5.1** | **12.5** | **4.5** | 22.0 | 5.5 | 17.0 | 4.7 | 21.1 | 5.0 |
| 7.0 | 12.2 | 5.1 | 8.7 | 4.5 | 19.5 | 5.5 | **13.0** | **4.7** | **16.6** | **5.0** |
| 8.0 | **9.4** | **5.1** | 7.4 | 4.5 | 14.5 | 5.5 | 10.6 | 4.7 | 12.4 | 5.0 |
| 9.0 | **8.2** | **5.1** | 6.6 | 4.5 | **12.3** | **5.5** | 9.3 | 4.7 | 10.2 | 5.0 |
| 10.0 | 7.2 | 5.1 | 6.0 | 4.5 | 10.1 | 5.4 | **8.2** | **4.7** | **8.9** | **5.0** |
| 11.0 | 6.8 | 5.0 | **5.7** | **4.5** | 8.7 | 5.4 | 7.5 | 4.6 | 8.0 | 5.0 |
| 12.0 | | | 5.5 | 4.5 | 7.8 | 5.4 | **7.1** | **4.6** | 7.0 | 5.0 |
| 13.0 | | | 5.3 | 4.5 | **7.2** | **5.4** | 6.6 | 4.6 | **6.3** | **5.0** |
| 14.0 | | | 5.2 | 4.5 | 6.6 | 5.4 | 6.4 | 4.6 | 5.9 | 5.0 |
| 15.0 | | | 5.1 | 4.5 | 6.1 | 5.4 | | | 5.6 | 5.0 |
| 16.0 | | | **5.0** | **4.5** | 5.7 | 5.4 | | | **5.5** | **4.9** |
| 17.0 | | | 4.8 | 4.5 | **5.4** | **5.4** | | | 5.3 | 4.9 |
| 18.0 | | | 4.8 | 4.5 | 5.1 | 5.4 | | | 5.1 | 4.9 |
| 19.0 | | | 4.7 | 4.5 | 5.0 | 5.3 | | | 5.0 | 4.9 |
| 20.0 | | | 4.7 | 4.5 | 5.0 | 5.3 | | | 4.9 | 4.8 |
| 21.0 | | | **4.6** | **4.5** | **4.9** | **4.8** | | | | |
| 22.0 | | | 4.5 | 4.5 | 4.8 | 4.3 | | | | |
| 23.0 | | | 4.5 | 4.5 | 4.8 | 4.2 | | | | |
| 24.0 | | | 4.5 | 4.5 | 4.8 | 4.1 | | | | |
| 25.0 | | | 4.4 | 4.4 | 4.7 | 4.1 | | | | |
| 26.0 | | | **4.4** | **4.4** | 4.7 | 4.1 | | | | |
| 27.0 | | | 4.4 | 4.5 | | | | | | |
| 28.0 | | | 4.4 | 4.5 | | | | | | |
| 29.0 | | | 4.4 | 4.4 | | | | | | |
| 30.0 | | | 4.4 | 4.4 | | | | | | |

Exact sampling depths (metres): 626 (1, 2.75, 4.5, 6.25, 8, 9.75), 239 (1.5, 6.5, 11.5, 16.5, 21.5, 26.5), 224 (1.5, 5.5, 9.5, 13.5, 17.5, 21.5), 223 (1, 3.25, 5.5, 7.75, 10, 12.25), 373 (1, 4, 7, 10, 13, 16).

Table S4: Sequences assigned in blank samples

| | July 2017 | July 2017 | October 2017 | October 2017 | October 2018 | October 2018 | October 2018 | October 2018 | October 2018 | August 2018 | August 2018 | August 2018 | August 2018 | August 2018 |
|---|---|---|---|---|---|---|---|---|---|---|---|---|---|---|
| **Lake** | **239** | **373** | **239** | **373** | **626** | **373** | **223** | **224** | **239** | **223** | **224** | **239** | **373** | **626** |
| **Total fish ASVs in blank sample** | 49 | 83 | 95 | 203 | 120 | 197 | 57 | 82 | 117 | 160 | 71 | 178 | 548 | 89 |
| *Salvelinus namaycush* ASV | 11 | 13 | 8 | 34 | 57 | 77 | 18 | 16 | 11 | 47 | 15 | 2 | 187 | 55 |
| *Perca flavescens* ASV2 | 18 | 0 | 21 | 0 | 60 | 0 | 0 | 0 | 37 | 1 | 0 | 30 | 0 | 22 |
| *Perca flavescens* ASV11 | 0 | 0 | 0 | 0 | 0 | 0 | 0 | 0 | 14 | 0 | 0 | 19 | 0 | 0 |
| *Coregonus* ASV4 | 6 | 0 | 31 | 0 | 0 | 0 | 0 | 0 | 30 | 0 | 0 | 103 | 0 | 0 |
| *Cottus cognatus* ASV7 | 9 | 0 | 23 | 0 | 1 | 0 | 0 | 0 | 23 | 1 | 0 | 22 | 0 | 0 |
| *Catostomus commersonii* ASV3 | 0 | 14 | 3 | 30 | 0 | 39 | 5 | 3 | 0 | 19 | 9 | 0 | 49 | 8 |
| *Cottus cognatus* ASV5 | 3 | 0 | 0 | 1 | 0 | 20 | 20 | 5 | 0 | 56 | 27 | 0 | 20 | 4 |
| *Margariscus margarita* ASV9 | 0 | 14 | 3 | 23 | 1 | 32 | 6 | 0 | 0 | 9 | 2 | 0 | 54 | 0 |
| *Chrosomus neogaeus* ASV6 | 0 | 33 | 3 | 115 | 1 | 29 | 0 | 10 | 0 | 0 | 0 | 0 | 235 | 0 |
| *Pimephales promelas* ASV8 | 2 | 9 | 0 | 0 | 0 | 0 | 8 | 48 | 0 | 27 | 18 | 0 | 0 | 0 |
| *Esox lucius* ASV12 | 0 | 0 | 0 | 0 | 0 | 0 | 0 | 0 | 2 | 0 | 0 | 2 | 0 | 0 |
| *Lota lota* ASV 39 | 0 | 0 | 1 | 0 | 0 | 0 | 0 | 0 | 0 | 0 | 0 | 0 | 0 | 0 |
| *Notropis atherinoides* ASV 37 | 0 | 0 | 1 | 0 | 0 | 0 | 0 | 0 | 0 | 0 | 0 | 0 | 0 | 0 |
| *Etheostoma nigrum* ASV 51 | 0 | 0 | 1 | 0 | 0 | 0 | 0 | 0 | 0 | 0 | 0 | 0 | 0 | 0 |

Table S5: Summary of sequences retained at each stage of bioinformatic analysis

| **Bioinformatics step** | **Mean ± 95% confidence error (per sample)** |
|---|---|
| Initial sequence number | 94,013 ± 6,389 |
| Initial sequence quality score | 33.0 ± 0.23 |
| Adapter trim (Cutadapt) | 83,575 ± 5,984 |
| Merge reads (Pear) | 83,340 ± 5,993 |
| Trim of remaining reads (Cutadapt) | 83,521 ± 5,983 |
| Length filter | 76,734 ± 5,954 |
| **Denoising pipeline** | **Mean ± 95% confidence error (all samples)** |
| Total sequences after length filtering | 34,990,837 |
| Total unique sequences after dereplication | 1,891,102 |
| Number of amplicon sequencing variants (ASVs) | 1031 |
| Number of non-chimeric ASVs | 373 |
| Number of reads mapped onto ASVs | 34,489,629 of 34,990,837 = 98.6% |

Table S6: Candidate model structures to assess species Amplicon Sequence Variant (ASV) count as a function of sampling depth, lake state (stratified or isothermal), and fish species

All models were fitted using glmmTMB in R v3.6.2. All models included a log offset of total DNA sequence counts, which accounts for variation in library size between samples. Here, lake depth is encoded as a continuous variable. Lake state is encoded as a factor with two levels (stratified or isothermal).

| Fixed effects model structure | Random effects | Link function | ΔAIC | Notes |
|---|---|---|---|---|
| Lake state * Depth * Species | Lake identity | Negative binomial | 0 | |
| (Lake state + Depth + Species)^2 | Lake identity | Negative binomial | 92.7 | |
| Lake state + Depth * Species | Lake identity | Negative binomial | 120.9 | |
| Lake state * Species + Depth | Lake identity | Negative binomial | 145.6 | |
| Species + Lake state | Lake identity | Negative binomial | 206.7 | |
| Lake state + Depth + Species | Lake identity | Negative binomial | 207.1 | |
| Lake state * Depth + Species | Lake identity | Negative binomial | 208.9 | |
| Species + Depth | Lake identity | Negative binomial | 209.9 | |
| Lake state * Depth * Species | Lake identity | Poisson | NA | Non-convergence |

Guide to model notation

* indicates that main effects and all interactions between terms are included.

^2 indicates that main effects and all two-way interactions between terms are included.

+ indicates the inclusion of main effects only

Table S7: Average depths of lake trout during stratification and turnover

Median and inter-quartile range of depths of lake trout during stratification and turnover, measured by acoustic telemetry. Average measures for the month and week before, as well as the day of sampling are provided.

| | Month | | Week | | Day | |
|---|---|---|---|---|---|---|
| **Lake** | **Median depth** | **IQR** | **Median depth** | **IQR** | **Median depth** | **IQR** |
| **223 Stratification** | 7.74 | 1.71 | 8.00 | 1.27 | 8.16 | 1.04 |
| **223 Turnover** | 1.73 | 1.55 | 1.64 | 1.37 | 1.77 | 1.35 |
| **626 Stratification** | 8.00 | 1.87 | 7.66 | 1.70 | 7.42 | 1.58 |
| **626 Turnover** | 1.87 | 1.41 | 1.90 | 1.19 | 2.19 | 1.09 |
| **239 Stratification** | 8.73 | 4.12 | 7.24 | 1.86 | 7.24 | 1.96 |
| **239 Turnover** | 3.69 | 6.80 | 5.27 | 5.24 | 4.30 | 5.33 |
| **224 Stratification** | 11.32 | 3.78 | 12.09 | 3.69 | 11.94 | 4.12 |
| **224 Turnover** | 6.51 | 10.48 | 8.91 | 10.44 | 13.61 | 10.73 |
| **373 Stratification** | 11.90 | 1.78 | 11.74 | 1.63 | 11.86 | 1.53 |
| **373 Turnover** | 3.83 | 7.56 | 2.46 | 5.37 | 1.55 | 3.28 |

Table S8: Candidate model structures to assess lake trout Amplicon Sequence Variant (ASV) count as a function of proportion of telemetry detections and lake state (stratified or isothermal).

All models were fitted using glmmTMB in R v3.6.2. All models included a log offset of total DNA sequence counts, which accounts for variation in library size between samples. The telemetry variable was coded as the average detections either a month or a week prior to sampling, or the date of sampling, for lake trout at that interval depth. This was expressed as a function of the total amount of telemetry detections in the lake for that time period, as well as a function of depth interval size. Lake state is encoded as a factor with two levels (stratified or isothermal).

| **Fixed effects model structure** | **Random effects** | **Link function** | **ΔAIC** |
|---|---|---|---|
| Month telemetry * Lake state | Lake identity | Negative binomial | 0 |
| Week telemetry * Lake state | Lake identity | Negative binomial | 0.5 |
| Month telemetry + Lake state | Lake identity | Negative binomial | 1 |
| Day telemetry * Lake state | Lake identity | Negative binomial | 1.3 |
| Week telemetry + Lake state | Lake identity | Negative binomial | 1.3 |
| Day telemetry + Lake state | Lake identity | Negative binomial | 1.8 |
| Month telemetry | Lake identity | Negative binomial | 2.6 |
| Lake state | Lake identity | Negative binomial | 2.6 |
| Week telemetry | Lake identity | Negative binomial | 3.0 |
| Day telemetry | Lake identity | Negative binomial | 3.4 |
| Month telemetry * Lake state | Month telemetry/ Lake identity | Negative binomial | 4.0 |
| Day telemetry * Lake state | Day telemetry/ Lake identity | Negative binomial | 5.3 |
| Week telemetry * Lake state | Week telemetry/ Lake identity | Negative binomial | NA Non-convergence |

Guide to model notation

* indicates that main effects and all interactions between terms are included.

+ indicates the inclusion of main effects only

Figure S1: Numbers of taxonomically assigned Amplicon Sequencing Variants (ASVs) using a local copy of the NCBI database

Panel A shows the incidence of unique ASVs from all taxonomic groups. Panel B shows the same ASVs weighted by their abundance in the whole dataset. 95.1% of DNA sequences mapped back onto fish ASVs from IISD Experimental Lakes Area. Darker blue colour in the fish category refers to the detection of fish species represented in the mock community (as opposed to at IISD Experimental Lakes Area).

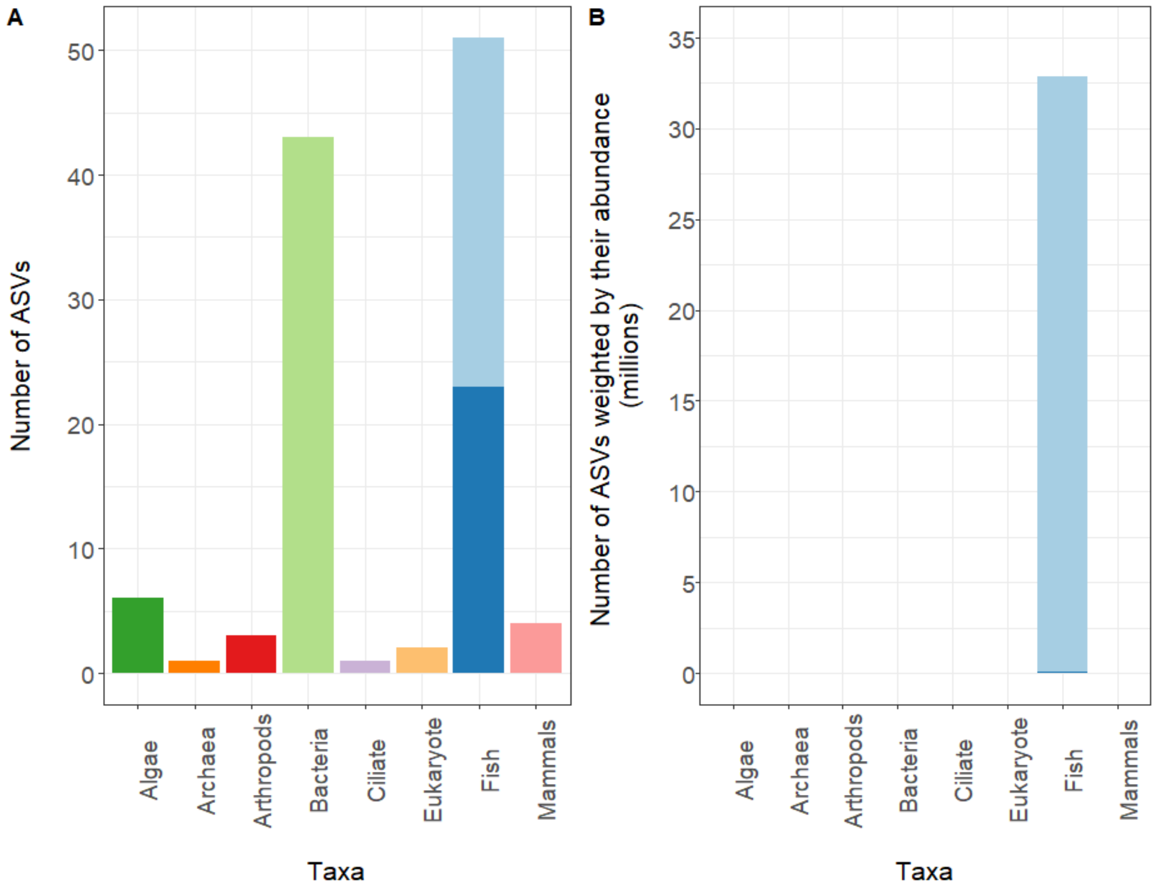
